# Fc-independent neutralization of SARS-CoV-2 by recombinant human monoclonal antibodies

**DOI:** 10.1101/2021.05.15.443978

**Authors:** Tal Noy-Porat, Avishay Edri, Ron Alcalay, Efi Makdasi, David Gur, Moshe Aftalion, Yentl Evgy, Adi Beth-Din, Yinon Levy, Eyal Epstein, Olga Radinsky, Ayelet Zauberman, Shirley Lazar, Shmuel Yitzhaki, Hadar Marcus, Angel Porgador, Ronit Rosenfeld, Ohad Mazor

## Abstract

The use of passively-administered neutralizing antibodies is a promising approach for the prevention and treatment of SARS-CoV-2 infection. Antibody-mediated protection may involve immune system recruitment through Fc-dependent activation of effector cells and the complement system. However, the role of Fc-mediated functions in the efficacious *in vivo* neutralization of SARS-CoV-2 is not yet clear. Delineating the role this process plays in antibody-mediated protection will have a great impact on the design of such therapeutics. Here, the Fc of two highly potent SARS-CoV-2 neutralizing human monoclonal antibodies, targeting distinct domains of the spike, was engineered to abrogate their Fc-dependent functions. The protective activity of these antibodies was tested against lethal SARS-CoV-2 infections in K18-hACE2 transgenic mice, both before or two days post-exposure in comparison to their original, Fc-active antibodies. Antibody treatment with both Fc-variants similarly rescued the mice from death, reduced viral load and prevented signs of morbidity. In addition, surviving animals developed a significant endogenous immune response towards the virus. Taken together, this work provides important insight regarding the contribution of Fc-effector functions in antibody-mediated protection, which should aid in future design of effective antibody-based therapies.

## Introduction

The ongoing COVID-19 pandemic caused by the severe acute respiratory syndrome coronavirus 2 (SARS-CoV-2) imposed a massive public health and economic crisis and continues to spread globally. Despite the onset of mass vaccination campaigns, the pandemic still exhibits unprecedented morbidity and mortality, highlighting the need for additional effective therapeutics. Human monoclonal antibodies (mAbs), specifically targeting viral surface proteins, have increasingly demonstrated prophylactic and therapeutic efficacy against various viruses including HIV, Ebola, the pathogenic beta-coronaviruses MERS-CoV and SARS-CoV and more recently for SARS-CoV-2 (*1-6*).

*In vitro* antibody-mediated viral neutralization is associated with steric interference with surface epitopes that are essential for virus entry into cells (as in case of antibodies targeting the receptor binding domain), membrane fusion, budding and more (*7*). However, *in vivo* antibody-mediated protection may further involve the immune system through Fc-dependent activation of effector cells and the complement system (*8-10*). The interplay between these two mechanisms and their individual contribution for effective viral neutralization is pathogen- and antibody-dependent. Indeed, while several studies demonstrated Fc-dependent activity for the neutralization of HIV, influenza and Ebola (*11-14*), other studies argued the opposite (*15*). It should be noted that for moderate or non-neutralizing antibodies *in vitro*, efficient Fc-activation of the immune system could augment and sometimes even be crucial for effective *in vivo* protection (*12, 13*). On the other hand, extensive activation of the immune system might also lead to antibody-dependent enhancement (ADE). Thus, it is essential to delineate the specific role and impact of antibody-based passive protection in the course of the disease and to design the Fc-mediated activity accordingly.

The use of passively-administered neutralizing antibodies is a promising approach for the prevention and treatment of SARS-CoV-2 infection (*16, 17*). Over the past year, many neutralizing antibodies were isolated, mainly targeting the spike protein and more specifically, its receptor binding domain (RBD) and N-terminal domain (NTD) (*4, 18-23*). Despite this body of work, the role of Fc-mediated functions in the efficacious *in vivo* neutralization of SARS-CoV-2, is not yet clear and only a limited number of studies have addressed this question so far. It was recently demonstrated that by enhancing the functionality of the Fc, the *in vivo* activity of a partially-neutralizing antibody, was significantly improved (*24*). Another set of studies have introduced loss-of-functions mutations into the Fc-region of SARS-CoV-2 neutralizing antibodies and evaluated their protective activity in infected animals, both as a prophylactic treatment and after infection. It was demonstrated that when given prophylactically, the protective activity of the mutated antibodies was minorly affected (*25, 26*). However, delaying the treatment, even for a short period (2-24 hours post infection) markedly reduced viral clearance and the protection conferred (*25-27*). The results of these studies may imply that once virus replication has begun, direct antibody neutralization is insufficient and other mechanisms that are based on Fc-activation are needed. Conversely, it can be argued that antibodies with superior neutralization activity, may be able to exert their direct activity even at late time points post infection, thus making Fc-dependent functions redundant.

As anti-SARS-CoV-2 antibody-based therapy is intended to treat COVID-19 patients or to be given prophylactically to immunocompromised populations, it is highly important to further investigate the Fc’s role in SARS-CoV-2 neutralization. Therefore, the role Fc activation plays will be a major influencing factor in therapeutic antibody design (*28*).

We have recently isolated a panel of SARS-CoV-2 neutralizing monoclonal antibodies from blood samples of severe COVID-19 patients (*22*). We were also the first to demonstrate that transgenic K18-hACE2 mice, infected with lethal doses of SARS-CoV-2, were fully protected by antibody administration as late as three days post infection (*4*), demonstrating the high potency of these antibodies. Here, we hypothesized that Fc-effector functions play a lesser role in the protective activity of highly potent SARS-CoV-2 neutralizing antibodies.

## Results

### Construction of Fc-engineered antibodies

Antibody-mediated activation of the Fc-gamma receptor (FcγR) plays a major role in viral neutralization *in vivo* (*7, 8*). The human FcγRI (CD64) has the higher affinity for both monomeric IgG and immune complexes, whereas FcγRIIa (CD32) and FcγRIIIa (CD16) strongly bind to IgG immune complexes (*10*). A single point mutation at N297 at the N-linked glycosylation motif Asn-X-Ser/Thr (N297/S298/T299) was previously reported to eliminate glycosylation and dramatically, but not completely, reduce binding to FcγR and to the complement system activator C1q (*29-32*). Maintaining the N-linked glycosylation while introducing double mutations at S298G/T299A was shown to specifically maintain binding to FcγRII, while abolishing binding to all other FcγR (*33*). Fully mutating the N-linked glycosylation motif by insertion of triple mutations at positions 297-299 inhibited this binding as well (*33*).

To study the role of Fc-mediated activation of immune effector functions in SARS-CoV-2 neutralization, we applied the human monoclonal antibody (mAb) MD65, recently reported to potently neutralize SARS-CoV-2 *in vitro* (*22*) and *in vivo* (*4*). The MD65 antibody backbone includes the triple mutation M252Y/S254T/T256E (YTE) in the Fc region (schematically depicted in Fig. 1A) aimed at increasing the antibody affinity towards the human FcRn at acidic pH, therefore prolonging its serum half-life (*34-36*). In order to diminish its Fc-dependent functions, MD65-YTE was further engineered to include the additional triple mutation N297G/S298G/T299A (MD65-AG-YTE, Fig. 1A). The engineered antibody was expressed in CHO cells. We first wished to confirm that its potency toward SARS-CoV-2 was retained. Indeed, comparison of the two versions of the MD65 (YTE and AG-YTE) confirmed that overall, their spike-binding performance is comparable (Fig. 1B), exhibiting apparent K_D_ of 0.4 nM versus 0.5 nM, for the YTE and the YTE-AG versions, respectively). Similarly, the two MD65 formats were evaluated by plaque reduction neutralization test (PRNT) and were shown to possess equivalent SARS-CoV-2 neutralization potency *in vitro* (NT_50_ of ∼40 ng/ml, Fig. 1C).

**Fig. 1.**
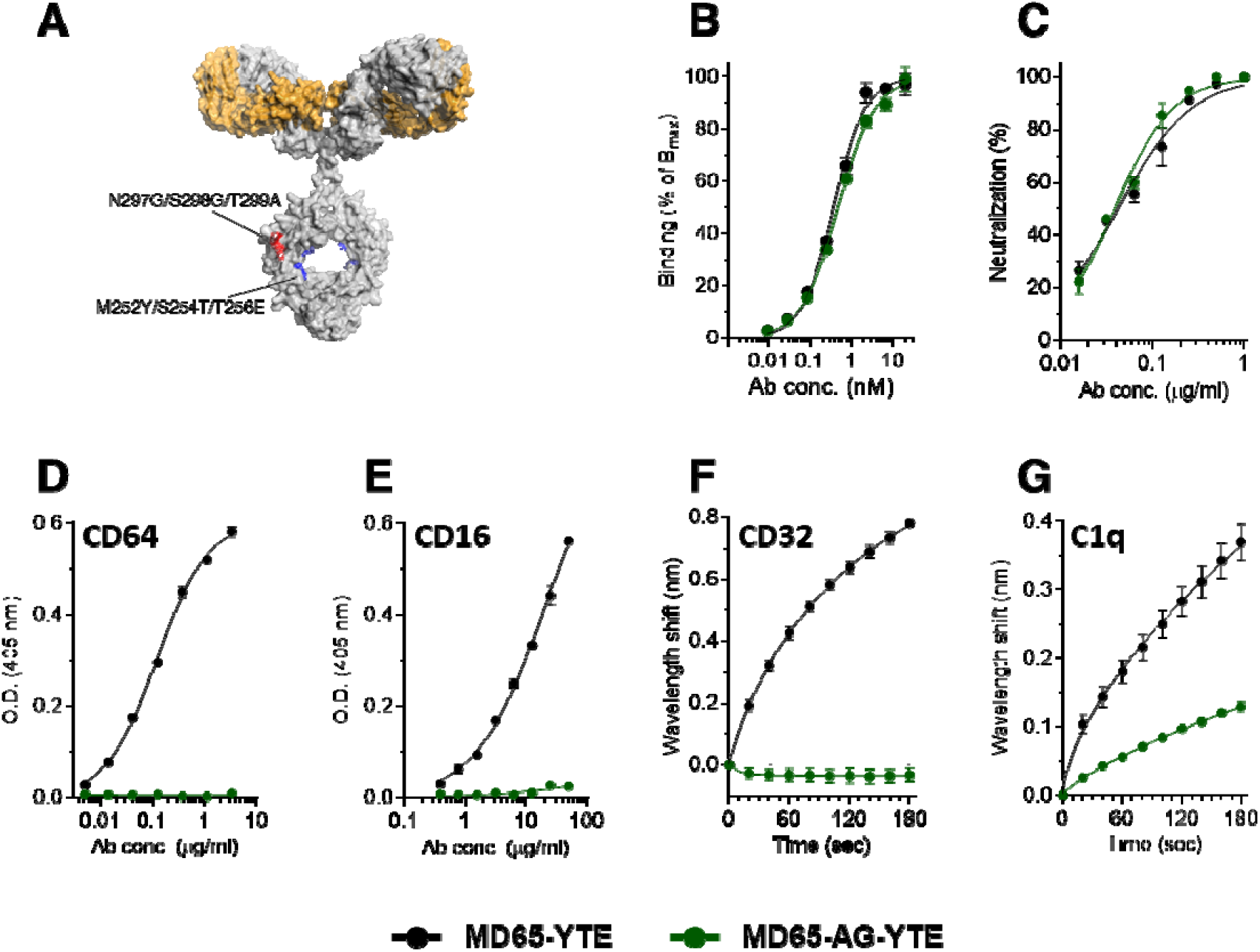
Binding characteristics of the MD65 antibody engineered versions. (**A**) Schematic representation of the mutations inserted into the YTE version (Red) and the AG-YTE version (Red and Blue). (**B**) Binding to immobilized SARS-CoV-2 spike protein, tested by ELISA. (**C**) *In vitro* neutralization of SARS-CoV-2 using plaque reduction neutralization test (PRNT). (**D, E**) Binding to immobilized CD64 (**D**) or CD16 (**E**), evaluated by ELISA. (**F**) BLI measurements of the binding of each antibody to immobilized CD32. (**G**) BLI measurements of the binding of C1q to the immobilized antibodies. Values are average ± SEM of triplicates of representative experiment (**B-E**) or of three independent experiments (**F, G**).

We next sought to confirm that the mutated MD65-AG-YTE antibody could not interact with the main Fc-mediated effector molecules. To this end, the binding of the antibodies to CD64 was evaluated using ELISA. As expected, MD65-YTE exhibited a dose dependent binding pattern whereas no binding of the AG-YTE variant could be detected in all tested concentrations (Fig. 1D). Similarly, while MD65-YTE showed robust binding to immobilized CD16, no interaction was observed with the MD65-AG-YTE antibody (Fig. 1E). The binding to CD32a was evaluated using biolayer interferometry (BLI), where the receptor was immobilized to the sensor and then reacted with either version of MD65. Indeed, while MD65-YTE induced significant wavelength shift indicating binding to the receptor, no binding was observed for the MD65-AG-YTE antibody (Fig. 1F). It was also of interest to determine whether the incorporation of these mutations had also affected the antibody ability to interact with the complement system. Since the complement cascade is initiated by the binding of C1q to the IgG-antigen immune complex, we evaluated its ability to interact with the MD65 antibody variants. Each antibody was immobilized on a BLI sensor, followed by incubation with a constant concentration of C1q which upon binding to the antibodies, induce a measurable wavelength shift. It was found that C1q could efficiently bind MD65-YTE, whereas its binding to the AG-YTE variant was greatly impaired (Fig. 1G). Non-linear fitting of the binding curves revealed that the affinity and the extrapolated Bmax values of C1q toward AG-YTE were about 260 and 50 times lower, respectively, when compared to the binding values toward the YTE version. These results suggest that the MD65-AG-YTE has retained only residual binding capability to C1q and completely lost its ability to bind to the Fc-receptor family.

It has been profoundly established that a major aspect of the monoclonal antibody therapeutic effect, is based on NK cell mediated antibody-dependent cellular cytotoxicity (ADCC) (*7*). Therefore, we have conducted a series of *ex vivo* functional assays to assess ADCC induced by both MD65-YTE and MD65-AG-YTE using enriched primary NK cells (pNK). A model system was tailored to specifically inspect the mAb-pNK cell activation axis (expression of CD107a and secretion of IFNγ and TNFα) (*37*). In order to induce Ag-mAb complex formation, tested mAbs were incubated with a reciprocal concentration gradient of SARS-CoV-2 spike and inert control protein of similar molecular weight, keeping total protein concentration constant. The membrane-associated CD107a staining assay indicates the extent of pNK cells’ activation and degranulation. As expected, only incubation of pNK cells with spike complexed with MD65-YTE induced elevated cell surface expression of CD107a (Fig. 2A). Accordingly, pNK cells secreted significantly less IFNγ and TNFα when incubated with a MD65-AG-YTE complexed with spike as compared to incubation with MD65-YTE-spike complex (Fig. 2B, C).

**Fig. 2.**
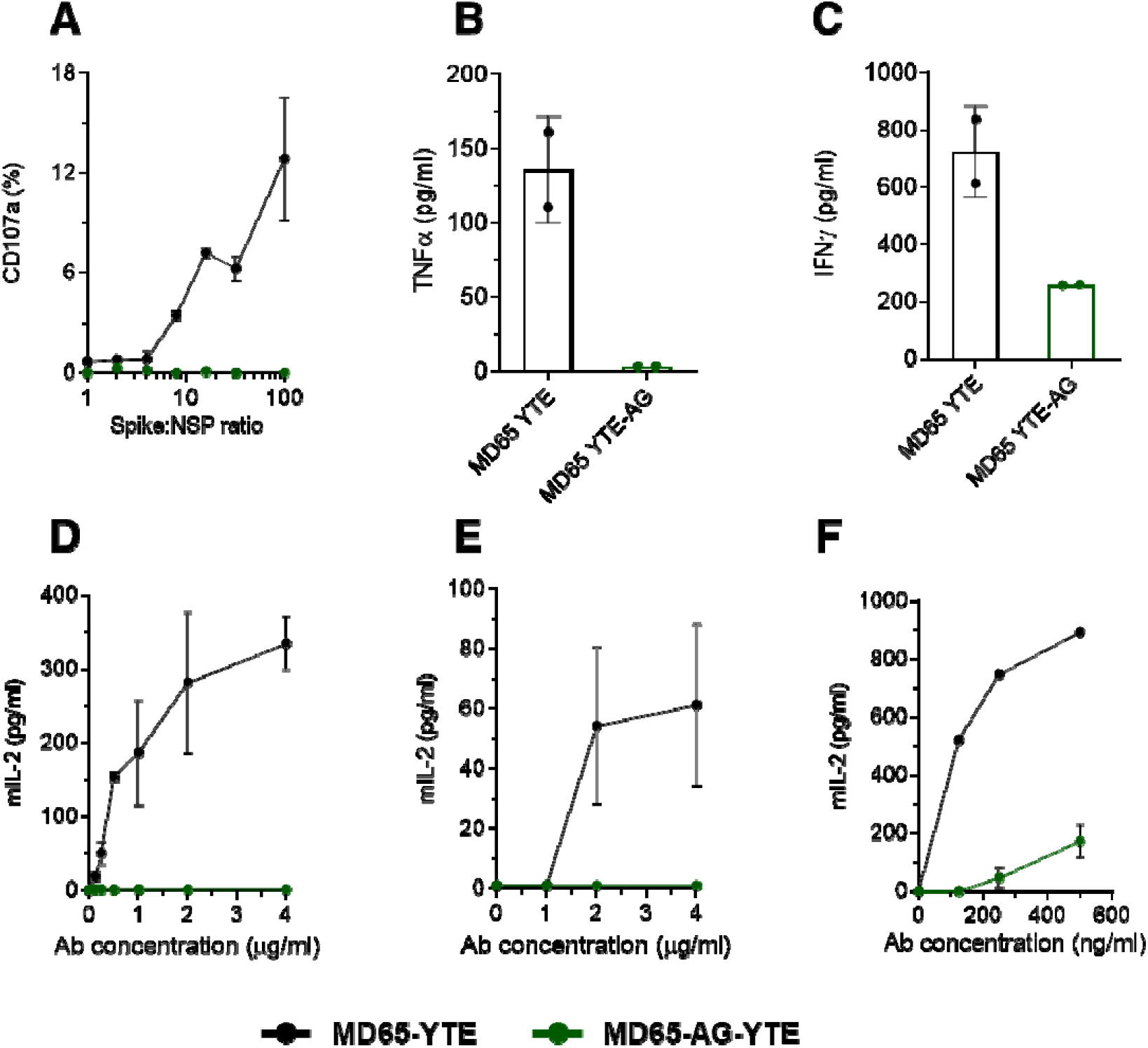
FcγR -mediated activation of cultured cells by the engineered versions of the MD65 mAb. (**A**) Primary NK cells were incubated for 4.5 hours with increasing concentrations of SARS-CoV-2 spike mixed with non-specific protein (NSP) that were complexed with either MD65-YTE or MD65-AG-YTE. Cell-surface expressed CD107a was evaluated by FACS. (**B**) TNFα or (**C**) IFNγ were measured in the supernatant of the cells that were incubated with spike only. BW5147 thymoma cells expressing human Fcγ receptors (**D**) CD16, (**E**) CD32 or (**F**) CD64 were incubated with spike complexed with antibodies for 16 hours. Secreted mIL-2 levels were measured by ELISA. Values are averages ± SEM of duplicates.

To further validate our findings, we assessed the activation potency of both MD65-YTE and MD65-AG-YTE that is mediated through Fcγ receptors. We have utilized the recently developed cell-based reporter system, designed to quantitate CD16 (FcγRIIIa), CD32 (FcγRIIa) and CD64 (FcγRI) mediated activation potency (*38*). Plates were coated with SARS-CoV-2 spike, incubated with increasing concentrations of either MD65-YTE or MD65-AG-YTE and washed to remove antibody excess. Reporter cells were then added to the immobilized Ag-mAb complexes and secreted mIL-2 was quantified, as a marker for reporter cell activation. Indeed, a distinct difference in activation potency between tested mAbs was observed, with MD65-YTE inducing significant, dose dependent levels of activation of CD16 and CD32 FcγRs (Fig. 2D, E). Similarly, MD65-YTE induced strong activation of CD64 presenting cells, while only marginal activation was caused by MD65-AG-YTE complexed with spike (Fig. 2F).

### Prophylactic activity of MD65-AG-YTE *in vivo*

After it was established that the AG-YTE version of MD65 antibody does not interact and activate the immune system via the Fc-dependent mechanisms, we sought to test its *in vivo* activity. The K18-hACE2 transgenic mice model was shown to faithfully recapitulate SARS-CoV-2 infections. Therefore, it serves as a reliable model to evaluate the efficacy of therapeutic strategies (*39-42*). We have recently applied this animal model and demonstrated that infection with a dose as low as 300 pfu of SARS-CoV-2 BavPat1/2020 strain, results in significant weight loss from day five after infection and death of 80% of the animals by day nine post-infection (*4, 43*). It was also shown that treatment of SARS-CoV-2 infected mice with MD65-YTE provided complete protection (*4*). Based on our previous experience, the bioavailability of MD65-YTE following IP administration is high, reaching plasma concentration peaks within 180 min and clearing with a half-life of about 4 days (*4*). It was thus important to assess whether the triple mutation incorporated into the Fc region of MD65-AG-YTE affect its pharmacokinetics parameters. As the clearance behavior of an antibody is a function of its affinity toward FcRn at low pH, we measured these parameters using BLI. Indeed, it was found that MD65-AG-YTE has retained the binding characteristics to FcRn (steady state K_D_ of 53 nM; Fig. S1).

Next, mice were intranasal (IN) infected with SARS-CoV-2 and concomitantly administered with 1 mg of either MD65-YTE or MD65-AG-YTE, intraperitoneal (IP). The clinical effect of the treatment of the two MD65 versions was then evaluated by measuring lung SARS-CoV-2 viral loads in infected mice six days post infection (the latest time point before mortality onset (*4*)). We employed qRT-PCR for the quantification of viral burden. In the control non-treated group, as expected, the virus propagated and reached very high concentrations (Fig. 3A). In contrast, treatment with either MD65-YTE or AG-YTE resulted in a significantly lower pulmonary viral loads by more than an order of magnitude compared to the untreated group (Fig. 3A). Further evaluation of the infectivity of the virus in these samples (by measuring its ability to infect VERO E6 cells), confirmed the qRT-PCR results and showed that treatment of infected mice with MD65 (either version) significantly inhibited (∼100-fold decrease) the ability of the virus to propagate with no apparent difference in effect between antibody version used (Fig. 3B). The severity of disease in the K18-hACE2 model is also evident as overexpression of pro-inflammatory cytokines (*44*). Based on the above results, it was speculated that the treatment will also reduce the serum levels of the pro-inflammatory cytokines. Thus, cytokines levels were measured in sera of mice infected with SARS-CoV-2 and either treated with MD65-YTE or MD65-AG-YTE, and compared to non-treated animals. The levels of the pro-inflammatory cytokines IL-1b and IL-6 were not affected by either antibody treatment (Fig. 3C, D). In contrast, mice that were treated with MD65-YTE exhibited significantly reduced levels of TNFα and IL-10, compared to untreated mice (Fig. 3E, F). Interestingly, while MD65-AG-YTE treated mice also displayed reduced levels of these two cytokines, this reduction was lesser than that observed for MD65-YTE-treated mice (Fig. 3E, F). One of the major roles of FcγR mediated activation of the immune system is to modify the expression of cytokines in response to infection (*7*). Accordingly, it is logical to assume that the incapability of MD65-AG-YTE to interact with FcγR is affecting the cytokine profile response. Yet, these changes are not essential for the antibody ability to induce efficient virus clearance.

**Fig. 3.**
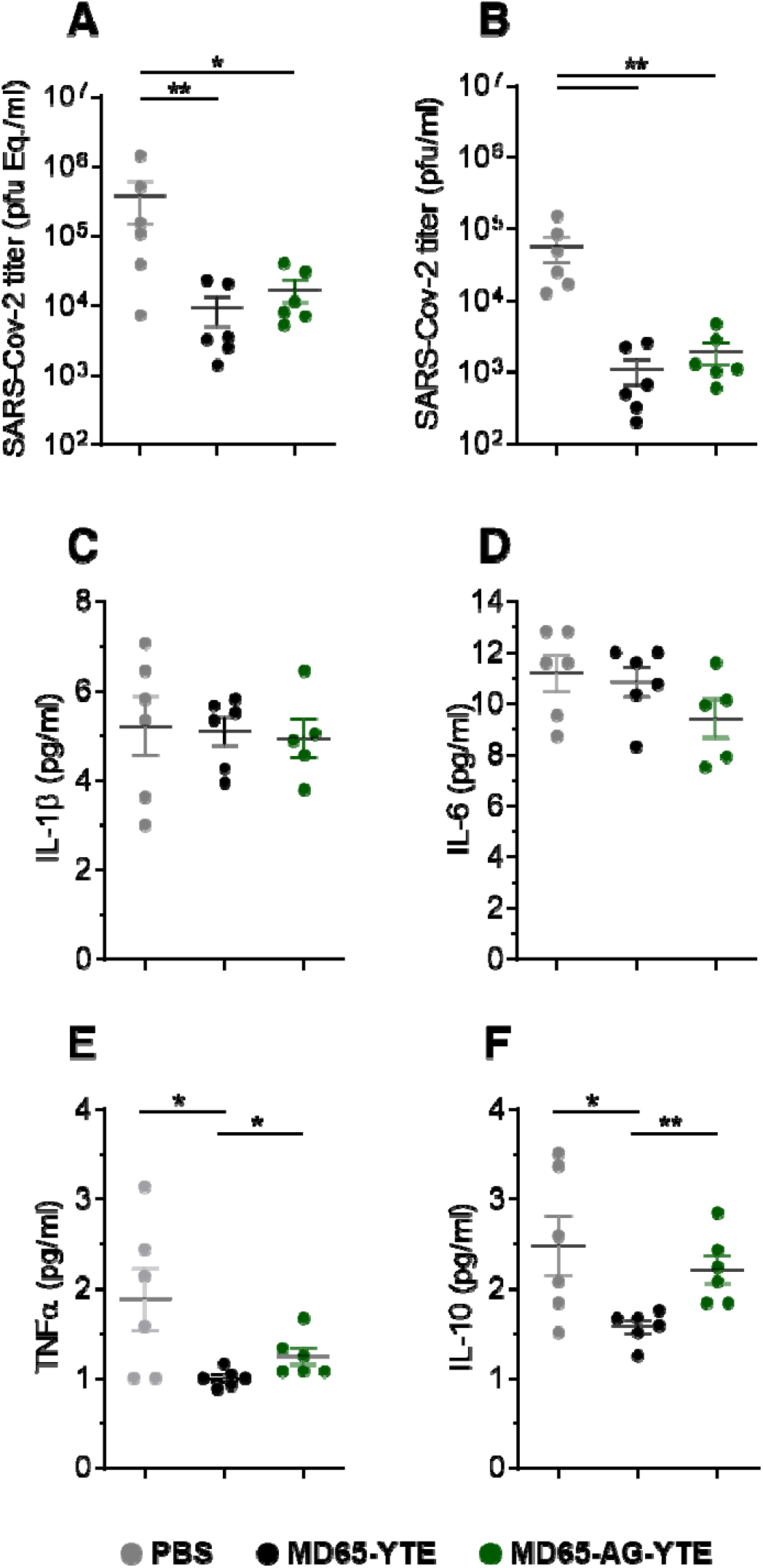
Viral load and cytokine levels following prophylactic administration of MD65 mAb versions. Lung and sera samples were collected six dpi from infected mice treated with the indicated MD65 versions or with PBS as control (n=6 for each group). (**A**) Viral load was quantified by qRT-PCR and expressed as equivalents of PFU/ml. (**B**) Infectious viral load determined by plaque assay. Cytokines levels in the sera were measured by multiplex kit (C) IL-1β, (D) IL-6, (E) TNFα and (F) IL-10. Data represent individual values and mean ± SEM. Horizontal bars, indicate statistical significance between paired groups. Significance (p) values were calculated using two-tailed unpaired t-test (* = p<0.05; ** = p<0.01).

### Fc-independent post-exposure protection of SARS-CoV-2 infected mice

The observation that when given as prophylaxis, MD65 activity is Fc-independent might be a reflection of direct interactions with the virus, inhibiting its ability to propagate and disseminate. While similar results were shown for several anti-SARS-CoV-2 antibodies, they all failed to confer protection when given post-infection, enforcing the notion that they mainly play a role in neutralizing the first steps of infection, before virus proliferation and spread (*25, 26, 45*). Therefore, it was of high importance to evaluate the relevance of Fc-mediated activation in MD65-based therapy in post-exposure treatment of SARS-CoV-2 infected K18-hACE2. To this end, mice were infected with 300 pfu of the SARS-CoV-2 BavPat1/2020 strain, treated two days post infection (dpi) with 1 mg of either MD65-YTE, MD65-AG-YTE or vehicle only (PBS) and further monitored up to 21 dpi. As expected, 80% of the infected animals of the vehicle-treated group have succumbed within 7 to 11 dpi (Fig. 4A), with a mean time to death of 8.5 days. In contrast, treatment with either versions of the MD65 antibodies was highly efficient in blocking disease progression, completely preventing weight loss and, most notably, fully protecting infected mice from death (Fig. 4A, B).

**Fig. 4.**
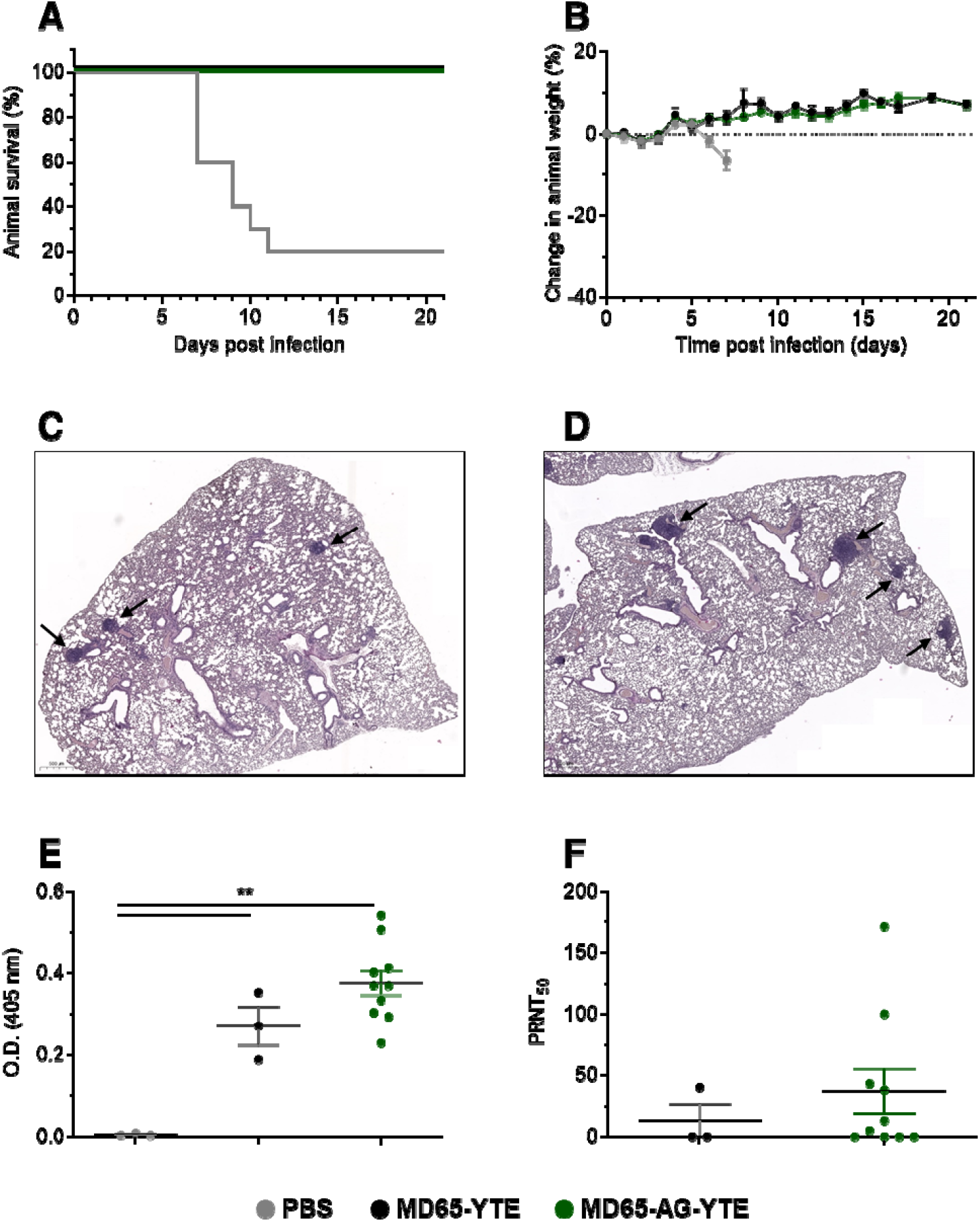
MD65 mAb-mediated post-exposure protection against SARS-CoV-2 infection of K18-hACE2 mice. Animals were intranasal infected with SARS-CoV-2 and treated two day later with 1 mg/mouse of MD65-YTE (n=3), MD65-AG-YTE (n=10) or with PBS as vehicle-control (n=10). **(A)** Kaplan-Meyer surviving curves. (**B**) Body weight profiles (data represents mean ± SEM). (**C-D**) Representative histological analysis of lung sections collected 21 dpi from MD65-AG-YTE (**C**) or MD65-YTE (**D**) treated mice. Black arrows indicate lymphoid aggregates. (**E-F**) Sera samples were collected at day 21 post-infection from the antibody-treated groups or from naïve mice and tested for the presence of endogenous (murine) antibodies against the SARS-CoV-2 spike by ELISA (**E**) and for *in vitro* neutralization capacity by PRNT (**F**). Data represent individual values and mean ± SEM. Horizontal bars indicate statistical significance. Significance (p) values were calculated using two-tailed unpaired t-test (** = p<0.01).

It was previously shown that SARS-CoV-2 infection of K18-hACE2 mice manifests as progressive and widespread viral pneumonia with perivascular and pan-alveolar inflammation, immune cell infiltration, edema, lung consolidation and distinctive vascular system injury that were apparent even three-weeks after infection (in surviving animals) (*41, 44, 46*). We have previously demonstrated that MD65-YTE based post-exposure treatment of SARS-CoV-2 infected K18-hACE2 mice, markedly prevented the formation of these pathological changes (*4*). Following the demonstration of the clinical efficacy of MD65 in treating infected mice, it was important to establish the pathological status of their lungs, especially in light of the fact that the activation of the immune system plays a major role in these changes. Accordingly, lung necropsy was performed 21 dpi in mice that were infected and treated as described with either MD65-YTE or MD65-AG-YTE. Histological evaluation demonstrated the lack of any major lung pathological changes or inflammation following treatment with either antibody (Fig. 4C, D). In accordance with our previous study, the only indication for a previous viral infection was the presence of scarce and well contained lymphoid aggregates, without any major differences between the MD65-YTE and the MD65-AG-YTE treated groups. The infiltration and accumulation of B-cells in the lungs of K18-hACE2 mice infected by SARS-CoV-2 was previously documented (*44*). The lymphoid aggregates observed in this study most probably contain B-cells that were recruited during initial accumulation of the virus in the lungs, within the first days after infection. Yet, the antibody treatment has halted the virus’ ability to further propagate and thus prevented the progression of inflammation. The observation that both antibody-treatments were equally effective, further substantiates the conclusion that efficient neutralization of SARS-CoV-2 *in vivo* is Fc-independent, even in situations where the virus has already begun propagation and dissemination.

### Anti SARS-CoV-2 seroconversion in antibody-treated animals

We have previously shown that surviving K18-hACE2 mice infected with SARS-CoV-2 and treated with MD65-YTE subsequently developed an endogenous immune response towards the virus (*4*). As mentioned before, FcγR-mediated activation of the immune system (by the formation of immune complexes, activation of antigen presenting cells and more) is one of the key factors contributing to the elicitation of antigen-specific antibody response (*7, 8*). Our findings that post-exposure treatment of infected mice with MD65-AG-YTE conferred full protection without any apparent long-term pathologies, prompted us to examine whether it had any effect on their ability to develop anti SARS-CoV-2 humoral response. Sera of K18-hACE2 mice treated with either MD65-YTE or AG-YTE two dpi, were collected 21 dpi and evaluated for anti-SARS-CoV-2 spike antibodies, by ELISA. As expected, MD65-YTE treated mice developed a significant (p<0.01) response when compared to naïve mice (Fig. 4E). Similarly, the MD65-AG-YTE treated mice have also developed strong and significant (p<0.001) endogenous humoral response toward the spike (Fig. 4F), comparable to that of the MD65-YTE group.

Although it is not in the direct scope of this work, it was of interest to determine whether the developed endogenous antibody response by these animals possess virus neutralization capabilities. First, we have validated that the sera samples do not contain any traces of the MD65 antibodies that may bias the interpretation of the neutralization data. Next, the SARS-CoV-2 neutralization potency of each sample was evaluated *in vitro*. Interestingly, half of the mice treated with MD65-AG-YTE developed neutralizing antibody responses (Fig. 4F), whereas only one MD65-YTE treated animal did so. Yet, the experimental group size was too small to confirm statistical significance to this observation.

### Dose-dependent therapeutic efficacy of MD65-AG-YTE

While the data so far indicated that efficient post-exposure protection of SARS-CoV-2 infected mice is Fc-independent, we asked whether Fc-immune activation may prove beneficial if treatment is given under sub-optimal conditions. One possible setup for such a challenge is to decrease the antibody:virus ratio. Thus, SARS-CoV-2 infected K18-hACE2 mice were treated two dpi with either 100 μg or 10 μg doses of antibodies. Treatment with 100 μg of MD65-YTE resulted in full protection of all treated mice, with no apparent signs of morbidity (Fig. 5A, B). In the parallel group, treated with MD65-YTE AG, 83% of the animals survived (Fig. 5A), with only one animal succumbing to infection. By monitoring animal weight as a surrogate marker for their overall clinical status, a short and transient decrease was observed around day eight in the MD65-YTE AG group (Fig. 5B). Yet, the treated animals quickly recovered and returned to their initial weight. These results fit well with the pharmacokinetics parameters of MD65 (*4*), where at day eight its concentration in the blood is reduced by more than 70%. At this time point, the *a priori* reduced dose together with antibody clearance and the inability to activate the immune system might combine to reduce treatment efficacy. Nevertheless, it had no significant effect on the treatment outcome, as the animals recovered and survived.

**Fig. 5.**
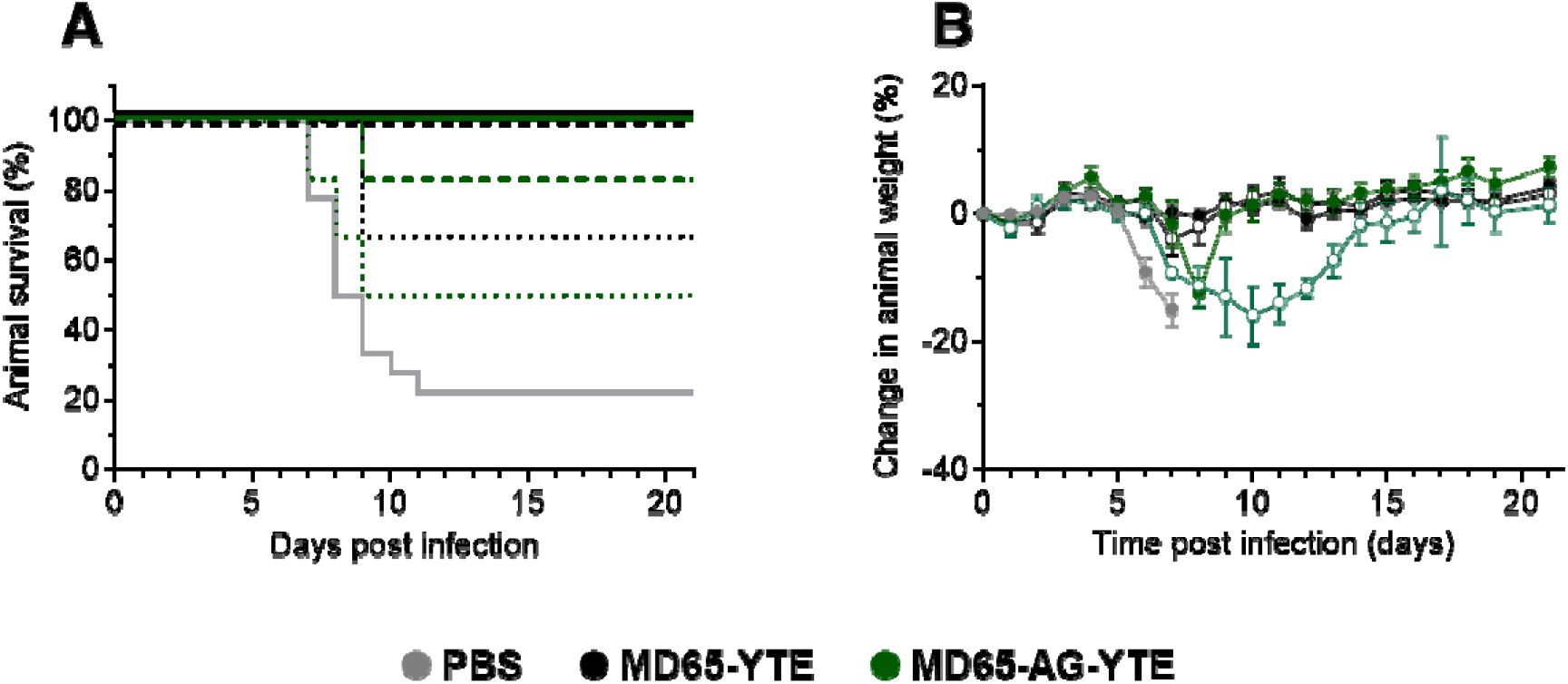
Dose-dependent post-exposure protection against SARS-CoV-2 infection of K18-hACE2 mice. Animals were intranasal infected with SARS-CoV-2 and treated two days later with either 100 μg/mouse (full lines) or 10 μg/mouse (dashed lines) of MD65-YTE (n=6 for each group), MD65-AG-YTE (n=6 for each group) or with PBS as control (n=8). **(A)** Kaplan-Meyer surviving curves. (**B**) Body weight profiles (empty and filled circles indicate treatment with 10 or 100 μg/mouse, respectively; data represents mean ± SEM).

Further reducing the antibody dose to 10 μg (100 times lower than the original dose) presented a similar trend. Treatment with either MD65-YTE or MD65-AG-YTE resulted in the survival of 67% and 50% of the animals, respectively (Fig. 5A). In the MD65-AG-YTE treated group, a marked reduction in the averaged animal’s weight was observed between days 7-14, at which time the surviving animals have fully recovered and returned to their initial weight (Fig. 5B). Accordingly, it may be concluded that even at sub-optimal treatment conditions, SARS-CoV-2 neutralization and protection is mostly Fc-independent. Yet, in the stage where antibody levels in the circulation are significantly low and the virus was not fully cleared, the recruitment of the immune system via Fc-activation will shorten the duration of the disease.

### Fc-independent post-exposure protection by anti-NTD antibody

In order to demonstrate that the potent Fc-independent protection against SARS-CoV-2 is not restricted to one selected antibody, we have chosen an additional anti-SARS-CoV-2 antibody, exhibiting a different neutralization mechanism. We and others have recently demonstrated that efficient neutralization of SARS-CoV-2 can be obtained by targeting the N-terminal domain (NTD), which does not directly interact with the host cell receptor (*23, 43*). The human-derived anti-NTD antibody BLN1 (*43, 47-53*) was expressed as either YTE or AG-YTE (using the same constant regions as those used for the MD65 antibodies). The binding of the two antibody versions was evaluated using ELISA against the spike protein, yielding similar pattern with an apparent K_D_ of 1.3 nM versus 0.7 nM, for the YTE and the AG-YTE versions, respectively (Fig. 6A). Accordingly, the two BLN1 formats exhibited comparable SARS-CoV-2 neutralization potency *in vitro*, with IC_50_ values of 20-40 ng/ml (Fig. 6B).

**Fig. 6.**
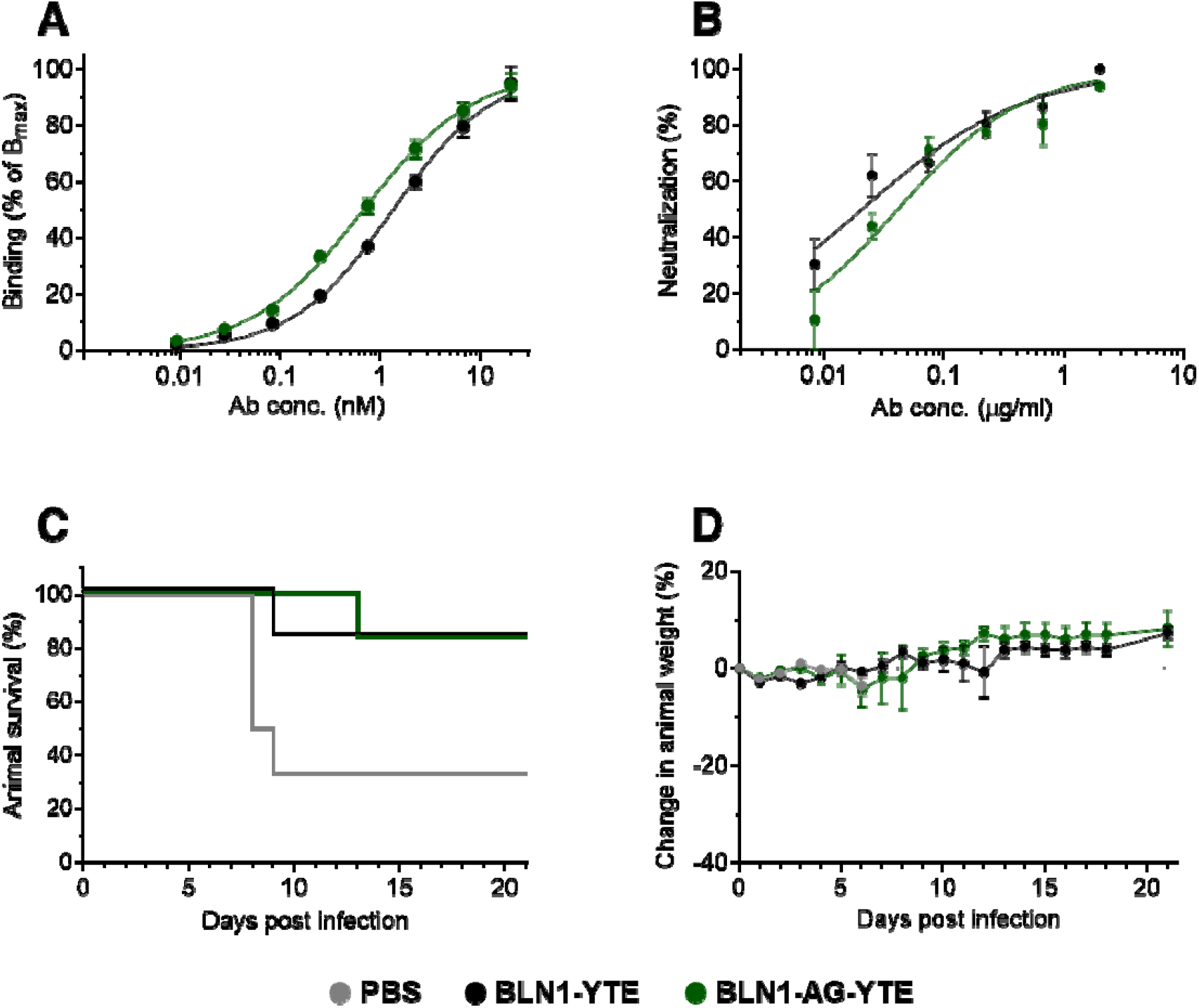
BLN1 mAb-mediated post-exposure protection against SARS-CoV-2 infection of K18-hACE2 mice. (**A**) Antibodies binding to immobilized SARS-CoV-2 spike protein. (**B**) *In vitro* neutralization of SARS-CoV-2, by PRNT. (**C, D**) Animals were intranasal infected with SARS-CoV-2 and two days later treated with 100 μg/mouse of BLN1 YTE (n=6), BLN1 AG-YTE (n=6) or with PBS as control (n=6). (C) Kaplan-Meyer surviving curves. (D) Body weight profiles (data represents mean ± SEM).

The neutralization efficacy of this antibody was further evaluated in the K18-hACE2 mice model. So far, the binding and the *in vitro* parameters of antibody BLN1 were highly similar to those determined for MD65. Thus, we decided to perform the *in vivo* experiment in the stringent format, initiating treatment two days post infection, and using the lower dose of 100 μg of either version of BLN1. Indeed, in this settings, treatment of the infected group with BLN1 YTE resulted in protection of 83% of the animals (Fig. 6C) with one death at day nine. The surviving animals showed no sign of illness or morbidity, as evident by weight monitoring (Fig. 6D). Similar results were obtained in the experimental group that was treated with BLN1 AG-YTE, where one animal succumbed on day 13, also resulting in 83% survival (Fig. 6C). It is also interesting to note that unlike the MD65-AG-YTE group, no weight loss (or any other clinical signs) was observed in the group that was treated with BLN1 AG-YTE (Fig. 6D). Taken together, these results suggest that efficient *in vivo* antibody-based neutralization of SARS-CoV-2 is Fc-independent, regardless of the specific molecular neutralization mechanisms.

## Discussion

Delineation of the role and impact of Fc-mediated activity in antibody-based passive protection against SARS-CoV-2 infection is of high interest and would assist in the design of such therapeutics. In this study, we have shown that two highly potent SARS-CoV-2 neutralizing antibodies do not rely on Fc-mediated immune activation to properly protect infected K18-hACE2 mice against lethal challenges.

At the aim of interfering with Fc-mediated effector functions, numerous mutations were examined, targeting amino acid positions that were found to interact with FcγR and the complement cascade (Lund et al. 1991; Sarmay et al. 1992; Idusogie et al. 2000; Oganesyan et al. 2008 Vafa et al. 2014; Wessels et al. 2016; Saunders 2019). However, since each of the proteins, involved in effector functions, interacts with the antibody’s Fc at slightly different positions, different sets of point mutations can either eliminate binding to a specific receptor or partially reduce binding to all receptors. One commonly used set of mutations creating an Fc-null antibody, is L234A/L235A (LALA mutations) (*54*). These mutations are directed at two points involved in the interaction between Fc and FcγR. The LALA mutations were shown to reduce ADCC activity (*55*) and dramatically reduce binding to FcγRI, IIa and IIIa (*54*). Nevertheless, introduction of the LALA mutations was demonstrated to allow minimal, but detectable, activity (*28*). Indeed, addition of a third mutation, P329G (LALA-PG), was needed to completely abrogate binding of antibodies to the FcγR (*56*). Here, we took an alternative approach to eliminating the activation of Fc-mediated functions, by inserting a triple mutation that eliminates Fc-glycosylation. The N-linked glycosylation of the Fc was shown to be involved in all types of effector functions, including antibody-dependent cellular cytotoxicity (ADCC), antibody-dependent cellular phagocytosis (ADCP) and cell-dependent cytotoxicity (CDC). It was therefore expected that elimination of glycosylation will abolish all such functions (*31*). In addition to type I FcγR, the Fc domain also interacts with type II FcR’s, which are C-type lectin receptors, CD209 (DC-SIGN) and CD23 (*57*). These receptors mainly bind sialylated-Fc domains of IgG antibodies, and are involved in regulation of inflammation and B-cell selection (*58*). Although not tested in the current study, it can be assumed that the absence of glycosylation also eliminates binding of the Fc domain by these receptors.

Despite the interest in understanding the role of Fc in *in vivo* neutralization of SARS-CoV-2, only limited studies have addressed this point so far. Two studies have performed a side-by-side comparison of the *in vivo* activity of a mutated Fc (LALA or LALA-PG) versus fully active antibody (*26, 27*). Both studies have found that treatment outcome, when given post-exposure (2-24 hours post infection), was markedly affected if Fc-mediated activity was abrogated. Two other studies have also evaluated the activity of similarly mutated Fc and found that if given post-exposure, their ability to protect animals from SARS CoV-2 infection, was inferior to prophylactic treatment (*25, 45*). Yet, these studies did not include a comparison with non-mutated antibodies, and thus the results could not be unequivocally attributed to the lack of Fc-activation. Based on their results in K18-hACE2 model, Winkler et al (*26*) suggested that once the virus has already begun to propagate and disseminate, effective protection requires Fc-mediated effector activity of the immune system. In contrast to the above-mentioned works, we have demonstrated that effective antibody-based therapy of SARS-CoV-2 infection in the stringent K18-hACE2 mice model can be achieved without the activation of the Fc-dependent effector functions. Moreover, this observation was confirmed by using two antibodies, each targeting and neutralizing SARS-CoV-2 by totally distinct mechanisms. One possible explanation for this apparent discrepancy, is that antibodies that possess superior neutralization activity, may be able to exert their direct activity even at delayed time points post infection, thus making Fc-dependent functions redundant. Direct comparison of antibodies’ activity, based upon published data is impossible as each laboratory uses different techniques and models to characterize the neutralization activity and potencies of their antibodies. Nevertheless, both MD65 and BLN1 belong to a unique set of antibodies that were shown to protect SARS-CoV-2 infected mice even when treatment was initiated 2-3 days post exposure (*4, 43*). Thus, it is suggested that they can be regarded as an “elite” set of neutralizing antibodies, for whom favorable clinical outcome does not necessitate the activation of Fc-mediated functions.

Only following treatment with very low antibody dosage, a slight beneficial effect of Fc-activation was observed, with mice treated with MD65-YTE not showing signs of illness while those treated with MD65-YTE-AG exhibited transient reduction in body weight from day 7-14. While the YTE mutations increased antibody affinity towards the human FcRn at acidic pH and prolonged its serum half-life in humans up to several months, it did not affect the pharmacokinetics in mouse sera. In fact, we have found that MD65-YTE has a serum half-life of about 3.5 days (*4*) in C57BL/6 mice (which share the same genetic background as K18-hACE2). Thus, the clinical signs observed here from day seven, correlate with a significant drop in antibody levels to about 20% of the initial values. Although not examined here, it can be speculated that if MD65-AG-YTE levels were kept constant for several days, then mice would not exhibit any signs of disease even if treated with such low and sub-optimal doses.

While in this study we have employed a mouse model to study the role of Fc in the treatment of COVID-19, the loss-of-function triple mutations (AG) that were inserted, were aimed to abrogate the activation of the human Fc-mediated effector functions. Nevertheless, it was previously shown that various human IgG isotypes differentially activate murine effector cells and that the human IgG1 is the most potent activator of ADCC and antibody-dependent cellular phagocytosis in the mouse (*59*). Thus, we believe that based on the clear overlap in the human IgG1 ability to activate murine Fc-effector functions, the K18-hACE2 infection model would reliably predict the behavior of Fc-engineered antibodies in humans.

To summarize, several studies have recently shown that anti-SARS-CoV-2 antibodies *in vivo* potency relies on Fc-effector functions for the efficient post-exposure treatment and that the activation of these pathways can augment their neutralizing potency. Here, we demonstrate for the first time, a proof-of-concept that “elite” antibodies do not necessarily require the modulation of the immune system. Since the activation of the immune system may be beneficial even for these antibodies and as there is no apparent antibody dependent enhancement of disease (ADE) in COVID-19 treatment, the ability to activate Fc-effector functions should be maintained. Nevertheless, as anti-SARS-CoV-2 antibody-based therapy is envisioned, among other scenarios, to treat COVID-19 immunocompromised patients, the specific role of Fc-activation should be characterize for each antibody before it enters the clinic.

## Methods

### Expression of proteins and antibodies

The previously developed MD65 monoclonal antibody (mAb), harboring the “YTE” Fc mutations (MD65-YTE) for increased half-life (*4*), was further engineered to obtain an a-glycosylated (AG) Fc format (MD65-AG-YTE). For that matter, additional three mutations were inserted to the Fc sequence, aiming at disrupting the N-linked glycosylation motif Asn-X-Ser/Thr (N297G/S298G/T299A). The presence of the desired mutations was confirmed by sequencing and the integrity and purity of the antibody were analyzed using SDS-PAGE. ExpiCHO-S cells (Thermoscientific, USA, Cat# A29127) were cultured at 37°C, 5% CO2 at 95% air atmosphere and used for expression of the recombinant antibodies, Which were then purified using HiTrap Protein-A column (GE healthcare, UK). The SARS-CoV-2 spike (S) stabilized soluble ectodomain and S1 subunit were produced as previously described (*22*).

### ELISA

Direct ELISA was performed against SARS-CoV-2 spike (*60*). Maxisorp 96-well microtiter plates (Nunc, Roskilde, Denmark) were coated overnight with 1 μg/ml of spike in NaHCO3 buffer (50 mM, pH 9.6), washed, and blocked with PBST at room temperature for 1 hour. The secondary antibodies: AP-conjugated Donkey anti-human IgG (Jackson ImmunoResearch, USA, Cat# 709-055-149, lot 130049) or AP-conjugated Donkey anti-mouse IgG (H+L) minimal cross (Jackson ImmunoResearch, USA, Cat# 715-055-150, lot 142717) were applied, followed by the addition of PNPP substrate (Sigma, Israel, Cat# N1891).

### Binding to recombinant FcγRs

Binding to FcγRs was tested using ELISA. CD16 (R&D Systems, USA, Cat #4325-FC-050) and CD64 (Sino Biological, China, Cat# 10256-H08S) were used to coat 96-well plate at 2 µg/ml. For CD16, plate was incubated with antibodies MD65-YTE and MD65-AG-YTE starting at 100 µg/ml with 2-fold serial dilutions. For CD64, antibodies were used starting at 10 µg/ml with serial dilutions. MD65-Fab, at the same concentrations, was used as negative control. Detection was carried out as described above. Binding to CD32 was tested using biolayer interferometry (BLI). Sensors were loaded with CD32 (10 µg/ml), followed by a wash, and then incubated with antibodies MD65-YTE or MD65-AG-YTE. MD65-Fab was used as negative control.

### Complement binding

Binding to the complement cascade was evaluated through measurement of antibodies binding to C1q protein, which represent the first step in the complement activation. Binding assays were carried out using the Octet system (ForteBio, USA, Version 8.1, 2015) that measures biolayer interferometry (BLI). All steps were performed at 30°C with shaking at 1500 rpm in a black 96-well plate containing 200 μl solution in each well. Fab_2_G sensors were loaded with MD65 mAbs (YTE or AG-YTE) at 10 µg/ml, followed by a wash. The sensors were then incubated with C1q native protein (250 nM; US Biologicals, USA, Cat# C0010-10D) for 180 s and then transferred to buffer-containing wells for another 180 s. Binding was measured as changes over time in light interference, after subtraction of parallel measurements from a sensor loaded with MD65-Fab as negative control. Sensograms were fitted with a 1:1 binding model using the Octet data analysis software 8.1 (Fortebio, USA, 2015).

### CD107a Degranulation assay

Cell culture plates were pre-coated with SARS-CoV-2 spike antigen (in PBS) and an inert control protein of similar molecular weight, in reciprocal concentrations. Total protein concentration was 2 µg/ml throughout the gradient while relative part of each protein varied (0:100, 1:99, 2:98, 4:96, 8:92, 16:84, 32:68, 100:0); Coating performed at 4°C overnight. Plates were then washed twice with PBS and tested mAbs were introduced in 2 µg/ml for 1 hour incubation in 4°C for Ag-mAb complex formation. Plates were then washed twice with PBS and primary NK cells (pNK) were introduced at a concentration of 2.5e5 cells/ml in 200 µl/well (5e4 cell/well) in complete SCGM media diluted 1:10 with complete RMPI media, final assay media contained 30 u/ml of a recombinant human IL-2 and 4 µg/ml allophycocyanin conjugated anti-CD107a. Cells were incubated in 37°C, 5% C0_2_ incubator for 4.5 hours. plates were then centrifuged at 300 g for 5 min, assay media was collected, and cells were stained using 4 µg/ml allophycocyanin conjugated anti-CD107a, and DAPI viability dye. Cells Analysis performed using Beckman Coulter CytoFLEX V5-B5-R3 Flow Cytometer.

### FCγ receptor potency assay

Cell culture plates were pre-coated with SARS-CoV-2 spike antigen in a concentration of 2 µg/ml at 4°C, overnight. Plates were then washed twice with PBS and tested mAbs were serially diluted and introduced to wells for Ag-mAb complex formation. Plates were incubated for 1 hour in 4°C, washed twice with PBS and human FCγ receptor (FcγRIIIa, FcγRIIa and FcγRI) expressing BW5147 thymoma cells were introduced to the Ag-mAb presenting wells during16 hours incubation (*38*). Plates were then centrifuged at 300 g for 5 min, assay media was collected and murine IL-2 was quantified by ELISA. Supernatant of activated cells were collected and specific cytokines were quantifies by ELISA. 96-well plates were pre-coated (in 0.1 M, Na_2_HPO; pH 9.0) at 4°C overnight with 70 µl/well of 1 µg/ml of a relevant capture mAb: purified anti-human IFN-γ (Clone:NIB42, BioLegend); Purified anti-human TNF-α (Clone:MAb1, BioLegend); or purified anti-mouse IL-2 (Clone: ES6-1A12, BioLegend). Plates were then incubated with blocking solution containing 10% FBS in PBST (0.05% Tween-20), washed and incubated with the collected supernatant for 2 hours, followed by addition of relevant detection mAbs: biotin anti-human IFN-γ (Clone: 4S.B3, BioLegend); biotin anti-human TNF-α (Clone: MP6-XT22, BioLegend); or biotin anti-mouse IL-2 (Clone: JES6-5H4, BioLegend) SA-HRP (Jackson immunoresearch, PA USA) and TMB (Dako, Denmark) used for detection of mIL-2.

### Plaque reduction neutralization test (PRNT)

Plaque reduction neutralization test (PRNT), performed essentially as described (REF) using SARS-CoV-2 (GISAID accession EPI_ISL_406862) strain that was kindly provided by Bundeswehr Institute of Microbiology, Munich, Germany and Vero E6 (ATCC® CRL-1586TM), obtained from the American Type Culture Collection. Half-maximum inhibitory concentration (IC_50_) was defined as mAb concentration at which the plaque number was reduced by 50%, compared to plaque number of the control (in the absence of mAb).

### Animal experiments

Treatment of animals was in accordance with regulations outlined in the U.S. Department of Agriculture (USDA) Animal Welfare Act and the conditions specified in the Guide for Care and Use of Laboratory Animals (National Institute of Health, 2011). Animal studies were approved by the local ethical committee on animal experiments (protocol number M-51-20). Female K18-hACE2 transgenic (B6.Cg-Tg(K18-ACE2)2Prlmn/J HEMI) were maintained at 20-22 °C and a relative humidity of 50±10% on a 12 hours light/dark cycle, fed with commercial rodent chow (Koffolk Inc.) and provided with tap water ad libitum. The age of the animals at the time of the onset of experiments ranged between9-14 weeks old. All animal experiments involving SARS-CoV-2 were conducted in a BSL3 facility. Infection experiments were carried out using SARS-CoV-2, isolate Human 2019-nCoV ex China strain BavPat1/2020 that was kindly provided by Prof. Dr. Christian Drosten (Charité, Berlin, Germany) through the European Virus Archive – Global (EVAg Ref-SKU: 026V-03883). The original viral isolate was amplified by 5 passages and quantified by plaque titration assay in Vero E6 cells, and stored at -80^°^C until use. The viral stock DNA sequence and coding capacity were confirmed as recently reported (*61*). SARS-CoV-2 BavPat1/2020 virus diluted in PBS supplemented with 2% FBS (Biological Industries, Israel) was used to infect animals by intranasal instillation of anesthetized mice. For mAbs protection evaluation, mice were treated IP either at the time of infection or 2 days post-infection. Control groups were administered with PBS at the indicated times. Body weight was monitored daily throughout the follow-up period post-infection. Mice were evaluated once a day for clinical signs of disease and dehydration. Euthanasia was applied only when the animals exhibited irreversible disease symptoms (rigidity, lack of any visible reaction to contact).

### Measurement of viral RNA by qRT-PCR

Viral load in lungs of SARS-CoV-2 infected mice was quantified by qRT-PCR and by plaque assay. Lungs were grinded in 1.5 ml PBS and 200 µl were added to LBF lysis buffer. RNA was extracted using RNAdvance Viral Kit on a Biomek i7 automated workstation (Beckman Coulter, IN), according to the manufacturer’s protocol. Each sample was eluted in 50 µl of RNase-free water. RT-PCR was performed using the SensiFASTTM Probe Lo-ROX One-Step kit (Bioline, UK). Primers and probe sequences, targeting the SARS-CoV-2 E gene, were based on the Berlin protocol published in the WHO recommendation for the detection of SARS-CoV-2 and as described before (*4*). The thermal cycling reaction was performed at 48°C for 20 min for reverse transcription, followed by 95°C for 2 min, and then 45 cycles of: 15 s at 94°C; 35 s at 60°C for the E gene amplification. Cycle Threshold (Ct) values were converted to PFU equivalents (PFU Eqv.), according to a calibration curve determined in parallel.

### Lung histology

Lungs were rapidly isolated, fixed in 4% PBS-buffered formaldehyde at room temperature for one week, followed by routine processing for paraffin embedding. Serial sections, 5 µm-thick, were cut and selected sections were stained with hematoxylin and eosin (H&E) and examined by light microscopy. Images were acquired using the Panoramic MIDI II slide scanner (3DHISTEC, Budapest, Hungary).

## Acknowledgments

We wish to express our gratitude to our colleagues Adva Mechaly, Tomer Israely, Sharon Melamed, Hagit Achdout, Yfat Yahalom-Ronen, Hadas Tamir, Emanuelle Mamroud, Shay Weiss, Itai Glinert, Theodor Chitlaru, Moshe Manzur, Yaron Vagima, Liat Bar-On, Noa Madar-Balakirski, Amir Rosner and Shmuel C. Shapira for fruitful discussions and support.

## Data and materials availability

Antibodies are available (by contacting Ohad Mazor from the Israel Institute for Biological Research) for research purposes only under an MTA, which allows the use of the antibodies for non-commercial purposes but not their disclosure to third parties.

## Author contributions

T.N-P., A.E., R.A., E.M., D.G., M.A., Y.E., A. B-D, Y.L., E.E., O.R., A.Z., S.L., S.Y., H.M., A. P., R.R., and O.M. designed, carried out and analyzed the data, added fruitful discussions and reviewed the manuscript. R.R. and O.M. supervised the project. All authors have approved the final manuscript.

